# Hexarelin exerts neuroprotective and antioxidant effects against hydrogen peroxide-induced toxicity through the modulation of MAPK and PI3K/Akt patways in Neuro-2A cells

**DOI:** 10.1101/2020.11.16.384321

**Authors:** Ramona Meanti, Laura Rizzi, Elena Bresciani, Laura Molteni, Vittorio Locatelli, Silvia Coco, Robert J. Omeljaniuk, Antonio Torsello

**Author notes:** Corresponding author (EB).

## Abstract

Hexarelin, a synthetic hexapeptide, protects cardiac and skeletal muscles by inhibiting apoptosis, both *in vitro* and *in vivo*. Moreover, evidence suggests that hexarelin could have important neuroprotective bioactivity.

Oxidative stress and the generation of free radicals has been implicated in the etiologies of several neurodegenerative diseases, including amyotrophic lateral sclerosis, Parkinson’s disease, Alzheimer’s disease, Huntington’s disease and multiple sclerosis. In addition to direct oxidative stress, exogenous hydrogen peroxide (H_2_O_2_) can penetrate biological membranes and enhance the formation of other reactive oxygen species.

The aim of this study was to examine the inhibitory influence of hexarelin on H_2_O_2_-induced apoptosis in Neuro-2A cells, a mouse neuroblastoma cell line. Our results indicate that H_2_O_2_ reduced the viability of Neuro-2A cells in a dose-related fashion. Furthermore, H_2_O_2_ induced significant changes in the morphology of Neuro-2A cells, reflected in the formation of apoptotic cell bodies, and an increase of nitric oxide (NO) production. Hexarelin effectively antagonized H_2_O_2_ oxidative damage to Neuro-2A cells as indicated by improved cell viability, normal morphology and reduced nitrite (NO_2_^−^) release. Hexarelin treatment of Neuro-2A cells also reduced mRNA levels of caspases−3 and −7 and those of the pro-apoptotic molecule Bax; by contrast, hexarelin treatment increased anti-apoptotic Bcl-2 mRNA levels. Hexarelin also reduced MAPKs phosphorylation induced by H_2_O_2_ and concurrently increased p-Akt protein expression.

In conclusion, our results identify several neuroprotective and anti-apoptotic effects of hexarelin. These properties suggest that further investigation of hexarelin as a neuroprotective agent in an investigational and therapeutic context are merited.

## Introduction

Growth hormone secretagogues (GHS) are a class of synthetic oligopeptides and non-peptidyl molecules with endocrine and extra-endocrine properties. GHS are preferentially recognized and bound by the ghrelin receptor, known as growth hormone secretagogue receptor type-1a (GHS-R1a) [1]. The GHS-R1a is predominantly found in the hypothalamus and pituitary gland where it mediates the release of growth hormone (GH). The GHS-R1a is also implicated in the regulation of gastrointestinal motility, as well as energy and glucose homeostasis [2].

In addition to its endocrine effects, GHS also target peripheral tissues; to illustrate, GHS improve muscle function in several pathological conditions by inhibiting the apoptosis pathway, reducing NO release and counteracting inflammation [3,4].

Among the GHS, hexarelin (a synthetic hexapeptide), binds not only to GHS-R1a but also to the CD36 receptor and manifests varied beneficial effects [5–7] in diseases associated with muscle wasting, chronic heart failure [8], excitotoxicity, neurological disorders, epilepsy and diabetes [9]. Despite the emerging biological importance of hexarelin, its signalling mechanisms have been only partially elucidated. Some studies have demonstrated that hexarelin modulates activation of different intracellular pathways, like mitogen-activated protein kinases (MAPKs) and phosphoinositide 3-kinase (PI3K)/protein kinase B (Akt) [10,11], and, thereby, could indirectly influence intracellular calcium (Ca^2+^) concentrations [12]. Furthermore, hexarelin protects cells *in vitro* from apoptosis by inhibiting nitric oxide (NO) synthesis and reactive oxygen species (ROS) release, modulating caspases activity as well as the expression of proteins belonging to the BCL-2 family [6,12–16].

Oxidative stress is involved in the progression of many neuronal disorders, such as Alzheimer’s disease, Parkinson’s disease, Huntington’s disease and amyotrophic lateral sclerosis, as well as in cancer, diabetes and aging [17–19].

Hydrogen peroxide (H_2_O_2_) is the most abundant ROS generated through oxidative stress in mitochondria [20]. H_2_O_2_ is formed by dismutation of superoxide radical anions catalysed by superoxide dismutase (SOD); it is freely soluble in aqueous solution and can easily penetrate biological membranes [21]. The ready diffusibility of H_2_O_2_ in intercellular fluids and the intracellular space can enhance mitochondria damage and potentiate the intrinsic apoptotic pathway [21–23].

The antioxidant effects of hexarelin have not yet been assessed. Therefore, the primary aims of this study are to characterize the effects of hexarelin on H_2_O_2_-induced oxidative stress and to explore its neuroprotective mechanisms of action in a mouse neuroblastoma cell line (Neuro-2A cells).

## Materials and methods

### Chemicals

Hexarelin was synthesized as previously described [5]. Dulbecco’s Modified Eagle’s Medium (DMEM)-high glucose, hydrogen peroxide (H_2_O_2_), 3-(4,5-dimethylthiazol-2yl)-2,5-diphenyl tetrazolium bromide (MTT) and Griess reagent were purchased from Sigma-Aldrich (St. Louis, MO, USA). Penicillin, streptomycin, L-glutamine, trypsin-EDTA, phosphate-buffer saline (PBS) and fetal bovine serum (FBS) were obtained from Euroclone (Pero, Milan, Italy). Prior to assay, hexarelin was dissolved in ultrapure water, and then diluted in *culture medium* to final working concentrations.

### Cell cultures

Neuro-2A murine neuroblastoma cells were cultured in DMEM-high glucose supplemented with 10% heat-inactivated FBS, 100 IU/ml penicillin, 100 μg/ml streptomycin and 2 mM L-glutamine under standard cell culture conditions (37°C, 5% CO_2_). Confluent cultures were washed with PBS, detached with trypsin-EDTA solution, and used for experiments.

In each experiment, Neuro-2A cells were incubated with H_2_O_2_ alone or the combination of 100 μM H_2_O_2_ and 1 μM hexarelin for 24 h.

At the end of each experiments, cell viability and nitrite (NO_2_^−^) release were measured. Cells were homogenized for the extraction of total RNA and for intracellular protein determinations. Caspase-3, caspase-7, Bax and Bcl-2 mRNA levels were measured by real-time RT-PCR, whereas protein levels of ERK 1/2, p-38 and Akt were made by western blot as described below.

### Cell viability

Neuro-2A cells were plated in 96-well culture plates at the density of 4 × 10^4^ cells/well for 24 h, at 37°C. The day after seeding, the cells were treated with increasing concentrations (50 - 200 μM) of H_2_O_2_ or with 100 μM H_2_O_2_ and hexarelin (1 μM). After 24 h of treatment, a 10 μl aliquot of 5 mg/ml MTT was added to each well and incubated at 37°C for 3 h. Then, the culture medium was removed and a 200 μl aliquot of acidified isopropanol was added in order to dissolve the formazan crystals. Absorbance was read at 570 nm using a multilabel spectrophotometer VICTOR^3^ (Perkin Elmer, Waltham, MA, USA). Cell viability of control cells was set to 100% and the relative absorbance of experimental groups were converted to relative percentages (relative absorbance of experimental group/relative absorbance of control) x 100= % of viable cells.

### Observations of morphological changes

Neuro-2A cells were seeded in 6-well culture plates at a density of 80 × 10^4^ cells/well and incubated at 37°C for 24 h. Cells were incubated with H_2_O_2_ (100 μM) in the absence or presence of hexarelin (1 μM) and photographed 24 h later using a Motic AE2000 Inverted Microscope (Motic, Hong Kong).

### Griess assay

NO production was evaluated measuring the nitrite (NO_2_^−^) content of culture media with the Griess reaction (G4410, Sigma-Aldrich). Neuro-2A cells were plated in 96-well culture plates and treated with 100 μM H_2_O_2_ and 1 μM hexarelin for 24 h. At the end of the treatment, 100 μl of medium were transferred to a new 96-well plate and were mixed to 100 μl of Griess reagent 1X. Absorbance was measured at 540 nm with a VICTOR^3^ spectrophotometer (Perkin Elmer, Waltham, MA, USA), after 15 minutes in the dark. A standard curve with varied concentrations of sodium nitrite was conducted in parallel and used for quantification.

### Real-time PCR analysis

In order to monitor the apoptosis pathway, Neuro-2A cells were plated in 24-well culture plates at a density of 2 × 10^5^ cells/well and incubated and treated at 37°C for 24 h according to specific protocols. Following treatment, Neuro-2A cells were washed with PBS and total RNA was extracted using EuroGOLD Trifast reagent (Euroclone), according to the manufacturer’s instructions and quantified using a Nanodrop ND-1000 spectrophotometer (Thermo Fisher Scientific, Waltham, MA, USA). Reverse transcription was performed using iScript™ cDNA Synthesis Kit (Bio-Rad, Hercules, CA, USA) to equal amounts (140 ng) of RNA. Amplification of cDNA (21 ng) was performed in a total volume of 20 μL of iTaq Universal Probes Supermix (Bio-Rad), using Real-Time QuantStudio7 Flex (Thermo Fisher Scientific). After 2 minutes at 50°C and 10 minutes at 94.5°C, 40 PCR cycles were performed using the following conditions: 15 seconds at 95°C and 1 minute at 60°C. Relative mRNA concentrations of the target genes were normalized to the corresponding β-actin internal control and calculated using the 2^−ΔΔCt^ method.

### Western blot

Neuro2A cells were plated in 6-well culture plates at a density of 8 × 10^5^ cells/well, incubated at 37°C for 24 h and then treated as previously described. Following treatment, cells were rinsed with ice-cold PBS and lysed in RIPA buffer (Cell Signalling Technology, Danvers, MA, USA), supplemented with a protease-inhibitor cocktail (Sigma-Aldrich), according to the manufacture’s protocol.

Total protein concentrations were determined using the Pierce BCA Protein Assay Kit (Thermo Fisher Scientific). Equal amounts of protein (20 μg) were heated at 95 °C for 10 min, loaded on precast 4-12% gradient gels (Invitrogen), separated by electrophoresis, and transferred to a polyvinylidene difluoride (PVDF) membrane (Thermo Fisher Scientific). Non-specific binding was blocked with 5% dried fat-free milk dissolved in phosphate-buffered saline (PBS) supplemented with 0.1% Tween-20 (PBS-T) for 1h at room temperature (RT). After washes in PBS-T, membranes were incubated with the primary antibody overnight at 4 °C (Anti-Phospho-p44/42 MAPK (Erk1/2) (Thr202/Tyr204) rabbit antibody, #9101, Cell Signaling Technology, 1:1000; anti-p44/42 MAPK (Erk1/2) rabbit antibody, #4695, Cell Signaling Technology, 1:1000; anti-Phospho-p38 MAPK (Thr180/Tyr182) rabbit antibody, #4511, Cell Signaling Technology, 1:1000; anti-p38 MAPK rabbit antibody, #9212, Cell Signaling Technology, 1:1000; anti-Phospho-Akt rabbit antibody, #4060, Cell Signaling Technology, 1:2000; anti-Akt rabbit antibody, #4685, Cell Signaling Technology, 1:1000; anti-actin rabbit antibody, #A2066, Sigma-Aldrich, 1:2500), and then with a peroxidase-coupled goat anti-rabbit IgG (#31460, Thermo Scientific, 1:5000) for 1 h at room temperature.

Signals were developed with the extra sensitive chemiluminescent substrate LiteAblot TURBO (Euroclone) and detected with a LAS imaging system. Image J software (National Institutes of Health, Bethesda, MD, USA) was used to quantify protein bands.

### Statistical analysis

Statistical analysis was performed using the program GraphPad Prism (GraphPad Software, La Jolla, CA, USA). Values are expressed as mean ± standard error of the mean (SEM). Experiments were independently replicated at least three times. Student’s *t*-test was used for comparisons between two groups. One-way ANOVA followed by Tukey’s t-test was used for multiple comparisons. A *p*-value of less than 0.05 was considered significant.

## Results

### Effects of hexarelin on H_2_O_2_-induced toxicity in Neuro2A cells

Neuro-2A cells were treated with various concentrations of H_2_O_2_ (50 – 200 μM) for 24 h in order to assess the toxicity of H_2_O_2_ and to identify the smaller concentration which significantly and reproducibly reduced cell viability. H_2_O_2_ reduced cell viability in a concentration-dependent manner (Fig 1A). As H_2_O_2_ at 100 μM significantly and reproducibly decreased cell viability (*p*<0.001) compared with vehicle-treated controls, it was used in subsequent experiments.

**Fig 1.**
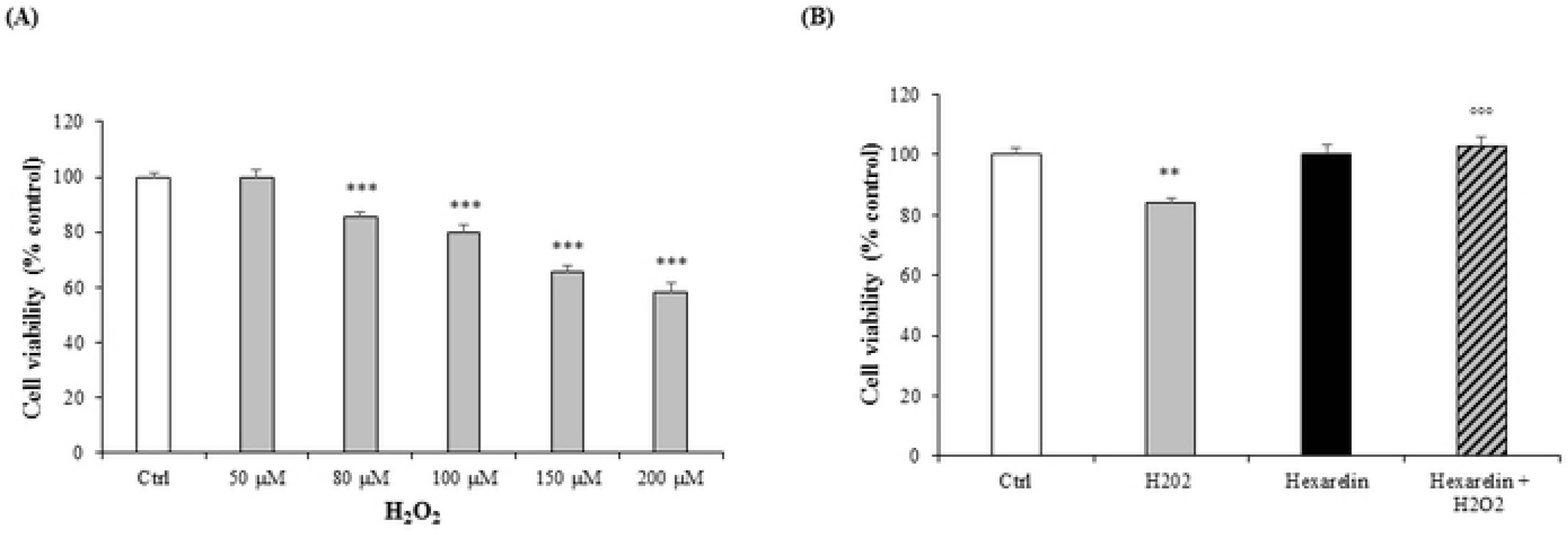
Cell viability in Neuro2A cells. Neuro-2A cells were treated for 24 h with different concentrations of H_2_O_2_ (0, 50, 80, 100, 150, 200 μM) and assessed for MTT assay (A). Neuro-2A cells were treated for 24 h with or without hexarelin (10^−6^ M) and H_2_O_2_ (100 μM) and assessed for MTT assay (B). All assays were performed in at least three independent experiments (*n=21*). Statistical significance: \*\**p*<0.01 vs CTRL, \*\*\**p*<0.001 vs CTRL; °°°*p*<0.001 vs H_2_O_2_.

Previous studies carried out in our laboratory demonstrated that exposure of Neuro-2A cells to various concentration of hexarelin (10^−8^ – 10^−5^ M) alone for 24 h did not reduce cell viability; consequently, 1 μM hexarelin was used in subsequent experiments (*data not shown*). The viability of cells treated with the combination of H_2_O_2_ and hexarelin for 24 h was similar to those of vehicle-treated controls but significantly (*p*<0.001) greater than those cells treated with H_2_O_2_ alone (Fig 1B).

### Morphological observations

The protective effects of hexarelin were confirmed by morphological observation using Motic AE2000 Inverted Microscope. The H_2_O_2_ treated Neuro-2A cells exhibited loss of confluence, disappearance of neurites, grouping and shrinkage of cells; these changes were even more pronounced at higher H_2_O_2_ concentrations (Fig 2A). These changes were obviously attenuated with hexarelin co-treatment (Fig 2B).

**Fig 2.**
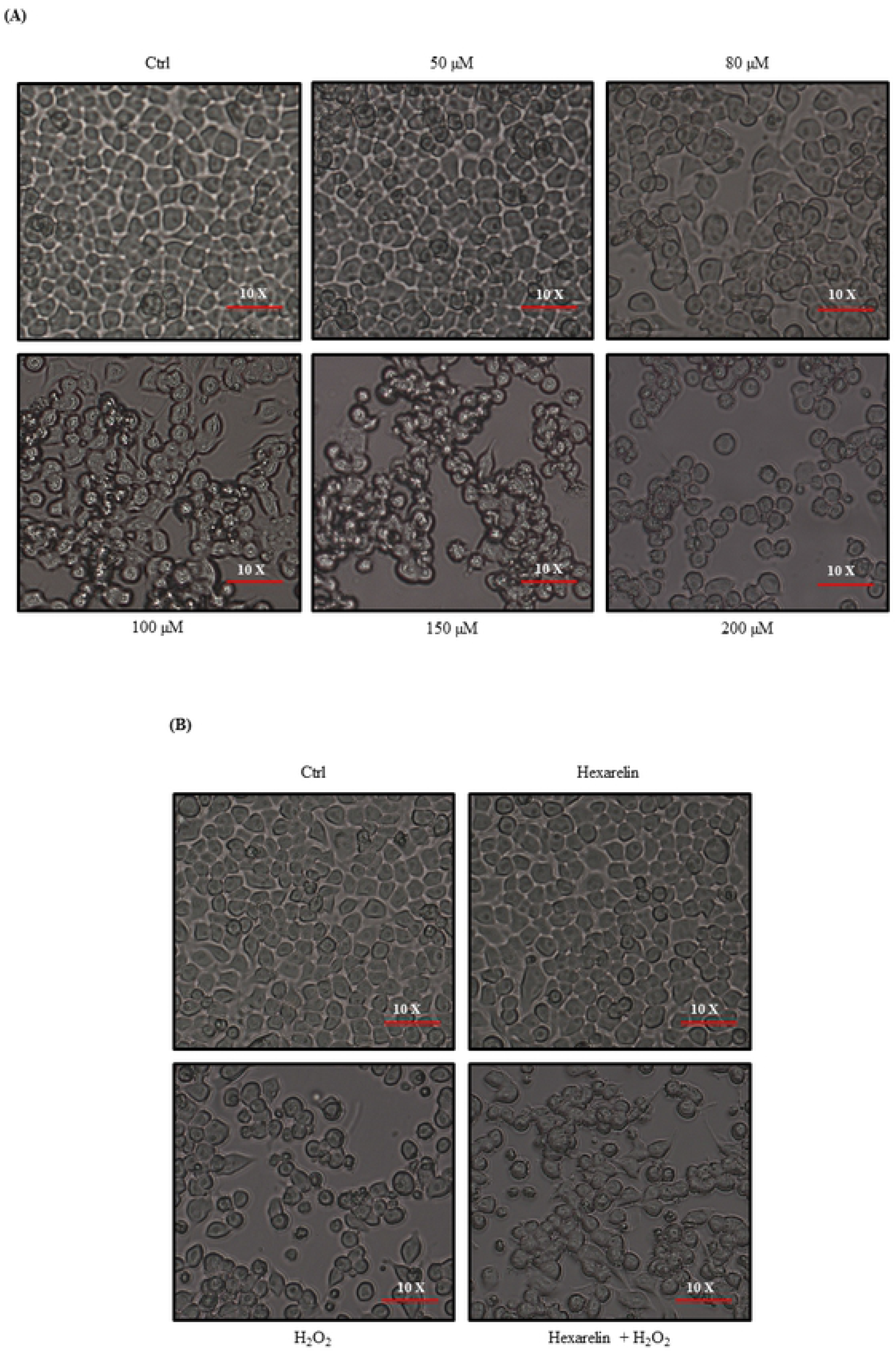
Morphological alterations in Neuro-2A cells. Morphological changes in Neuro-2A cells were observed by Motic AE2000 Inverted Microscope, at 10X magnification. Representative images of cells treated with increasing concentration of H_2_O_2_ (A) and in presence of hexarelin and/or 100 μM H_2_O_2_ (B).

### Effects of hexarelin on NO production on H_2_O_2_-induced Neuro2A cells

Nitric oxide (NO) is a highly reactive cytotoxic free radical, which can be induced by oxidative stress. To evaluate NO formation induced by H_2_O_2_, extracellular nitrite (NO_2_^−^) concentrations were measured by Griess assay.

Treatment of cells with 1 μM hexarelin alone had no significant effect on NO_2_^−^ release compared with vehicle-treated controls; by comparison, H_2_O_2_-treatment (100 μM) alone significantly increased levels of NO_2_^−^ (*p*<0.05) compared with vehicle-treated controls (Fig 3). In sharp contrast, extracellular NO_2_^−^ levels in cells co-incubated with 1 μM hexarelin and 100 μM H O were significantly (*p*<0.05) lower than in cells treated with H_2_O_2_ alone.

**Fig 3.**
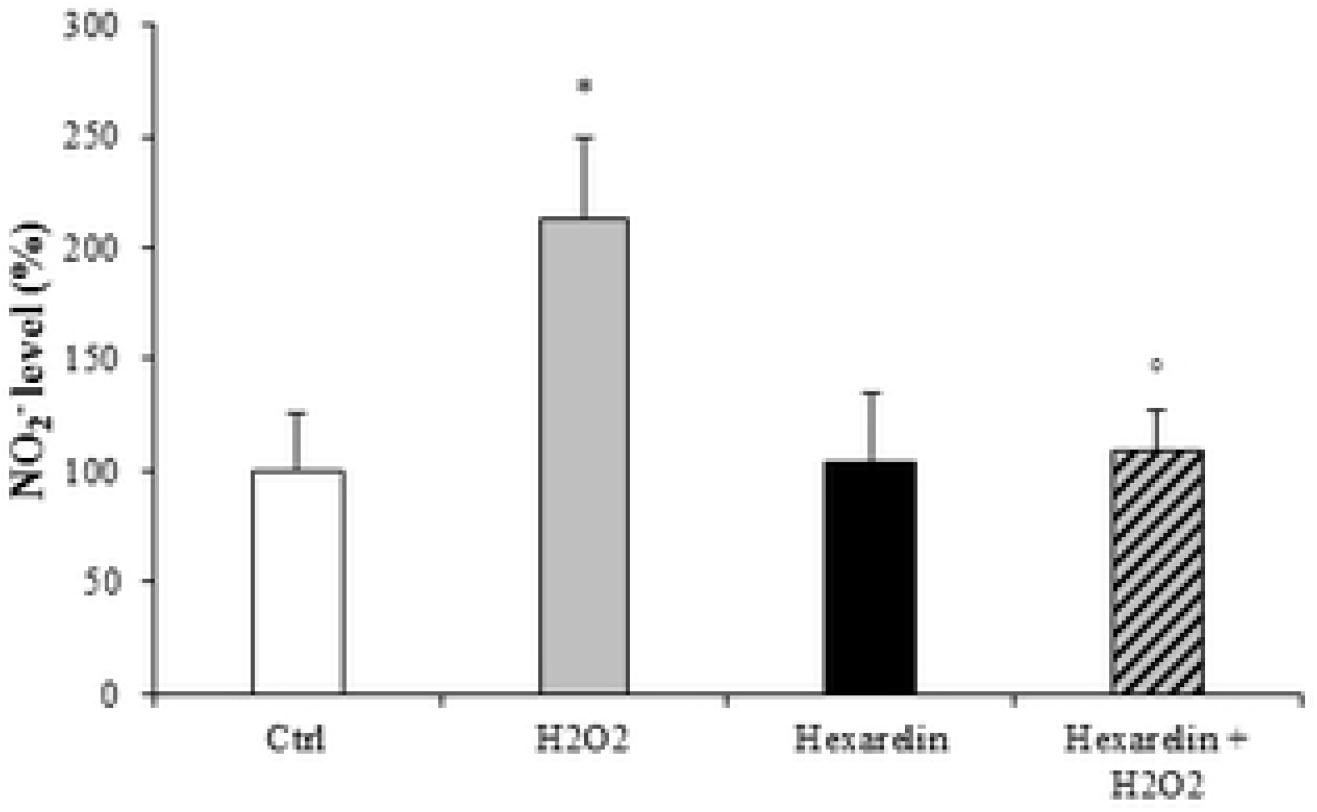
Effects of hexarelin on the extracellular NO_2_^−^ production. Neuro-2A cells were treated for 24 h with or without hexarelin and H_2_O_2_ 100 μM. The culture media was used for Griess reaction to measure NO_2_^−^ extracellular release. Data are expressed ad mean ± SEM of 4 replicates (*n=24*). Statistical significance: \**p*<0.05 vs CTRL; °*p*<0.05 vs H_2_O_2_.

These results suggest that the anti-apoptotic properties of hexarelin may be through an antioxidant mechanism.

### Effects of hexarelin on caspases – 3 and – 7 and on the modulation of apoptosis pathway

Caspase-3 and caspase-7 activation after oxidative damage induced by H_2_O_2_ occur at an early stage of apoptotic cell death; therefore, we hypothesised that hexarelin would inhibit caspase-3 and caspase-7 synthesis.

First, we demonstrated that mRNA expression of both caspases was increased by H_2_O_2_ in dose-dependent manners and that 100 μM H_2_O_2_ induced a significant increase of caspase-3 (*p*<0.01) and caspase-7 (*p*<0.001) mRNA expression (Figs 4A and 4B). Hexarelin did not affect mRNA levels of either caspases; by contrast, H_2_O_2_ significantly increased mRNA expression of both caspase −3 and −7 (*p*<0.001) (Figs 4C and 4D). Notably, the presence of 1 μM hexarelin in cells co-incubated with 100 μM H_2_O_2_ caused a non-significant reduction in caspase-7 mRNA (Fig 4D), but, significantly reduced caspase-3 mRNA (*p*<0.001, Fig 4C).

**Fig 4.**
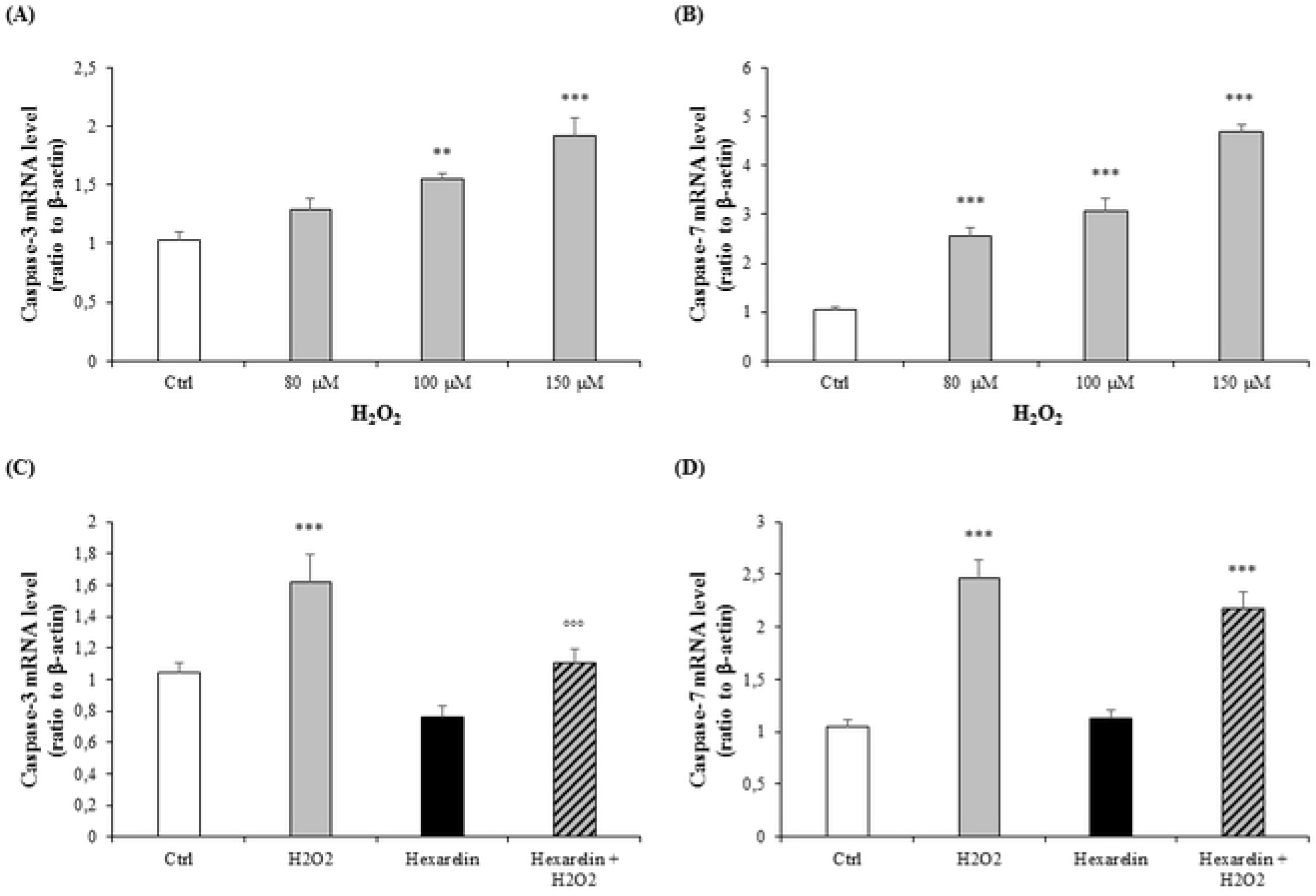
Analysis of caspase-3 and caspase-7 mRNA levels following H_2_O_2_ exposure and their modulation induced by hexarelin. Neuro-2A cells were treated with different concentration of H_2_O_2_ (80, 100, 150 μM) or co-incubated with hexarelin 10^−6^ M and H_2_O_2_ 100 μM for 24 h. Caspase-3 (A, C) and caspase-7 (B, D) mRNA levels were normalized for the respective β-actin mRNA levels. Data are expressed ad mean ± SEM of 3 replicates (*n=18*). \*\**p*<0.01 vs CTRL, \*\*\**p*<0.001 vs CTRL; °°°*p*<0.001 vs H_2_O_2_.

In addition to the activation of caspases, mitochondria also play a crucial role in the process of cell apoptosis with the activation of BCL-2 protein family. Increasing concentrations of H_2_O_2_ caused a trend toward an increase of the pro-apoptotic Bax levels (Fig 5A) and a trend toward a decrease of the anti-apoptotic Bcl-2 levels (Fig 5B). The co-treatment for 24 h of hexarelin and H_2_O_2_ in Neuro-2A cells showed the anti-apoptotic effect of hexarelin (Figs 5C and 5D). Hexarelin significantly reduced the mRNA expression of Bax induced by H_2_O_2_ (*p*<0.05) and increased expression of Bcl-2 mRNA (*p*<0.001).

**Fig 5.**
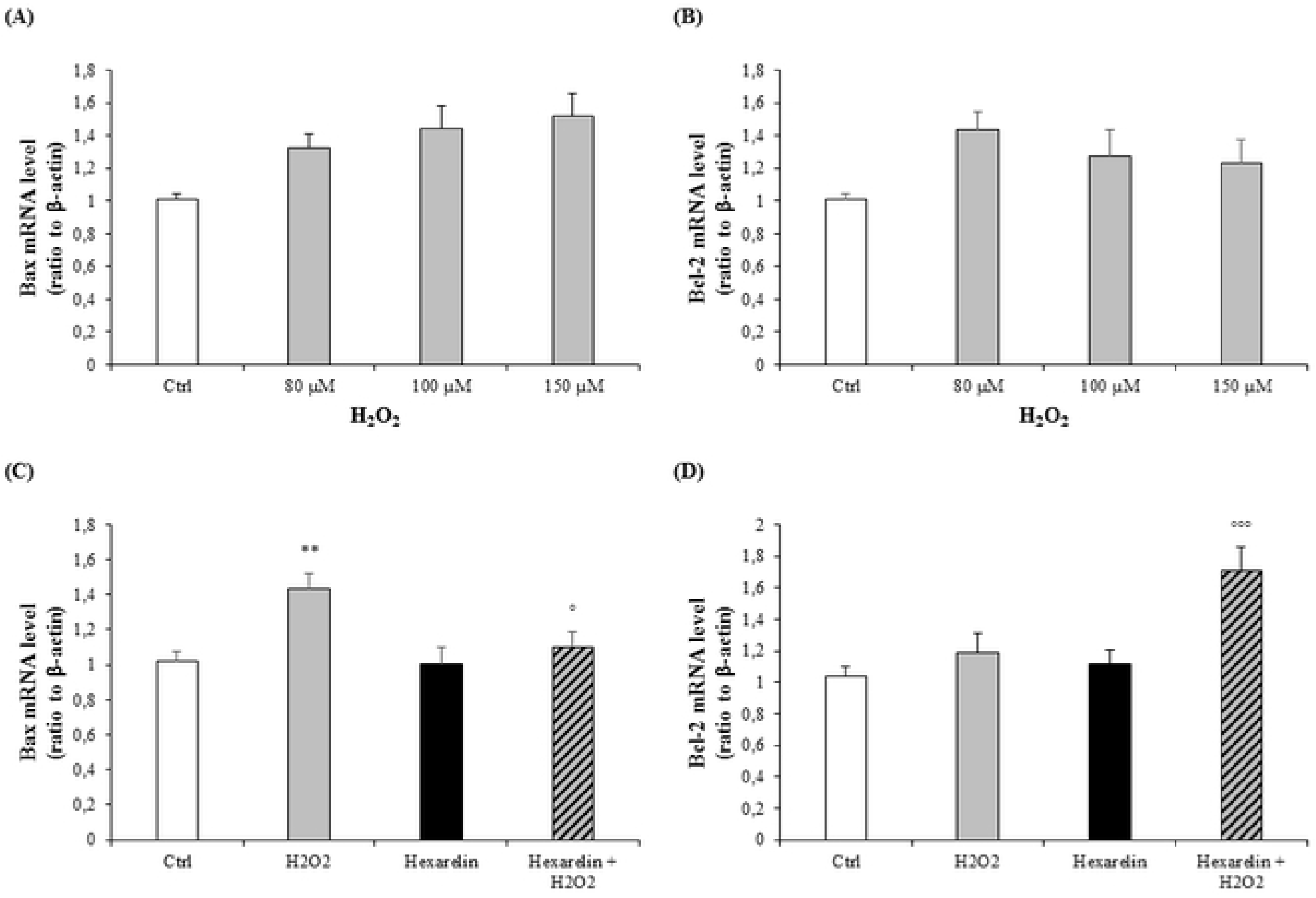
Analysis of mRNA levels of apoptosis markers following H_2_O_2_ exposure and their modulation induced by hexarelin. Neuro-2A cells were treated with different concentration of H_2_O_2_ (80, 100, 150 μM) or co-incubated with hexarelin 10^−6^ M and H_2_O_2_ 100 μM for 24 h. Bax (A, C) and Bcl-2 (B, D) mRNA levels were normalized for the respective β-actine mRNA levels. Data are expressed ad mean ± SEM of 3 replicates (*n=18*). \*\**p*<0.01 vs CTRL; °*p*<0.05 vs H_2_O_2_, °°°*p*<0.001 vs H_2_O_2_.

### Effects of hexarelin on ERK 1/2, p38 and Akt protein levels in H_2_O_2_- treated Neuro2A cells

As a consequence of the preceding findings, we hypothesized that hexarelin could modify MAPK signalling as well. Among members of the MAPK family, ERK and p38 are known to be associated with cell death or survival, respectively [10].

Exposure to 100 μM H_2_O_2_ significantly increased the p-ERK/t-ERK ratio (p<0.05), by comparison, 1 μM hexarelin did not affect ERK protein levels, both compared to the control group. Notably, co-incubation with hexarelin and H_2_O_2_ decreased p-ERK protein levels (Fig 6A).

**Fig 6.**
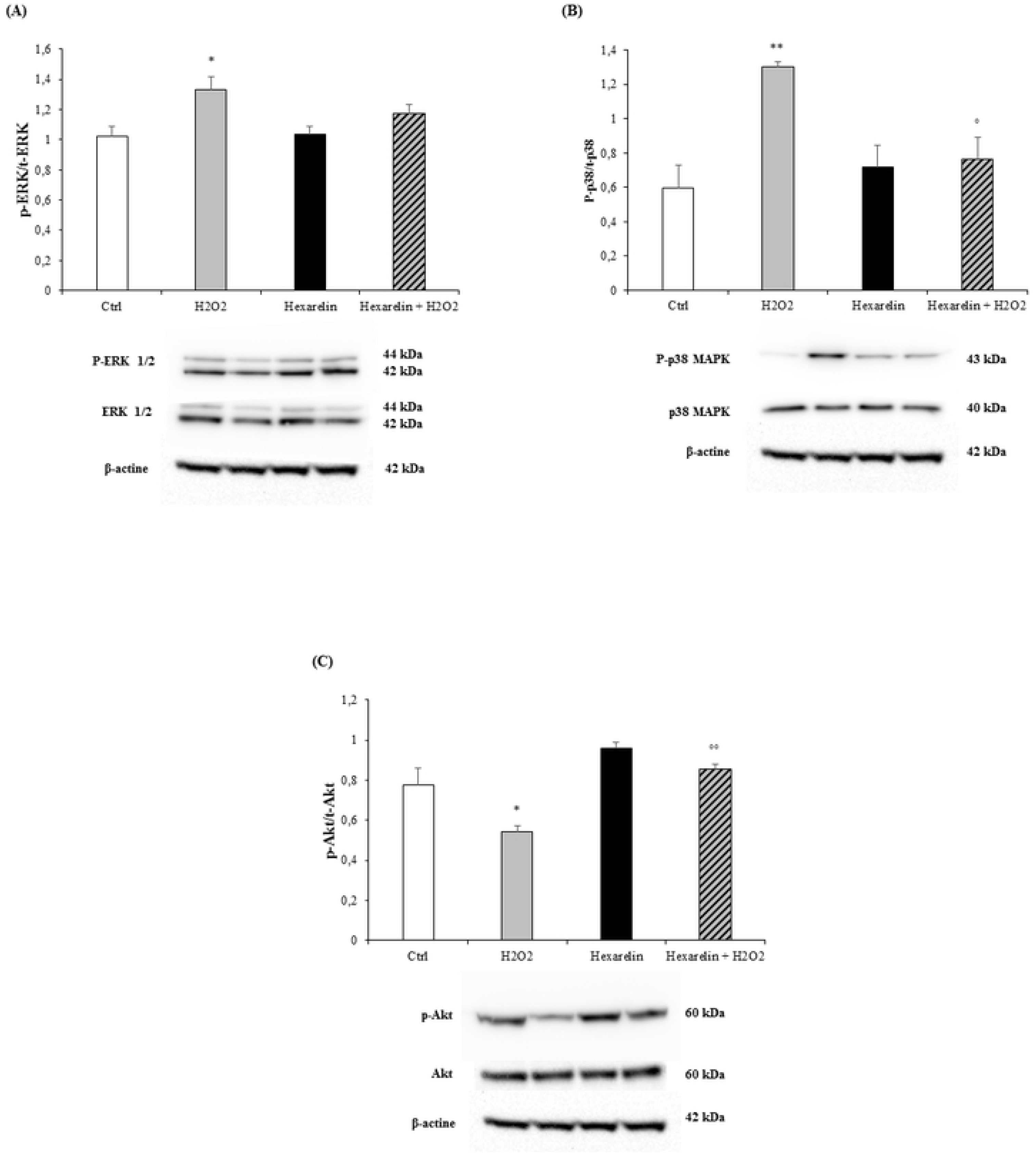
Hexarelin inhibits ERK and p38 activation and increase Akt activation after H_2_O_2_-stimulation. Neuro-2A cells were treated with or without hexarelin and H_2_O_2_ for 24 h and assessed in western blot for p-ERK/t-ERK (A), p-p38/t-p38 (B) and p-Akt/t-Akt (C). All assays were performed at least three independent experiments (*n=5*). Statistical significance: \**p*<0.05 vs CTRL; °°*p*<0.01 vs H_2_O_2_.

Western blot analysis revealed that hexarelin treatment of Neuro-2A cells exposed to H_2_O_2_ significantly decreased the level of p-p38 when compared to the H_2_O_2_-treated group (p<0.05). Hexarelin alone increased, non-significantly, p-p38 levels compared to controls while 100 μM H_2_O_2_ significantly increased p-p38/t-p38 ratios (p<0.01) (Fig 6B).

Furthermore, levels of p-Akt, associated with cell survival after oxidative stress [24], were not changed by hexarelin treatment alone compared to the control but were reduced by H_2_O_2_ treatment (*p*<0.05, Fig 6C). Notably, p-Akt levels in cells treated with hexarelin and H_2_O_2_ were significantly (*p*<0.01) greater than those in cells treated with H_2_O_2_ alone.

## Discussion

Our *in vitro* findings demonstrate that hexarelin protects mouse neuroblastoma cells from H_2_O_2_-induced damage.

Oxidative stress arises when the balance between oxidants and antioxidants is disrupted in favour of the former resulting in potential damage to the organism [21,24]. Although each neurodegenerative disease (NDD) has its own distinct etiology and differentially affects brain regions, NDDs share elements of oxidative stress, free radical generation and mitochondrial changes, leading to apoptosis [25].

Apoptosis is known to be one of the most sensitive biological markers for evaluating oxidative stress caused by imbalance between ROS generation and efficient operation of antioxidant systems [26,27]. Apoptotic cell death is an active process initiated by genetic programs and culminates in DNA fragmentation, characterized by morphological changes, including cell shrinkage, formation of membrane-packaged bits called apoptotic bodies [23], activation of caspases, nucleases, the inactivation of nuclear repair polymerases [28] and finally condensation of nuclei [29].

There are numerous inducers of oxidative stress that are able to cause cytotoxicity in *in vitro* models. In our study, we used H_2_O_2_ as it is an established method for measurement of potential neuroprotective antioxidants [30,31].

In this study H_2_O_2_ induced neuronal cytotoxicity in Neuro-2A cells in a dose-dependent manner. In order to determine the protective effect of GHS against H_2_O_2_-induced cytotoxicity, Neuro-2A cells were treated with hexarelin at 1 μM. The cell viability of the hexarelin treated group was similar to untreated control cells; by contrast, H_2_O_2_-induced cell death was significantly attenuated by hexarelin.

The protective effects exerted by hexarelin on H_2_O_2_-induced cytotoxicity determined by MTT assay was supported by morphological observations. Thus, evidence indicates Neuro-2A cells die as the result of apoptosis after H_2_O_2_ insult in dose-dependent manner, and that treatment with hexarelin attenuates the 100 μM H_2_O_2_-induced neuronal damage.

The protective effect of hexarelin was further confirmed by Griess assay. Excessive levels of NO, an important mediator of cellular communication at basal levels and implicated in the pathogenesis of NDDs [32], could be quantified by the measurement of extracellular NO_2_^−^, a primary stable products of NO breakdown. Previous studies report that H_2_O_2_-induced oxidative stress results in increased production of NO in neuronal and glial cells [33,34] through the induction of inducible nitric oxide synthase. In this study, we demonstrated that in Neuro-2A cells H_2_O_2_ increased extracellular NO_2_^−^ release and that cells treated with hexarelin showed lower NO_2_^−^ levels compared to the H_2_O-exposed cell group thereby demonstrating its protective effect against H_2_O_2_-induced oxidative stress.

Hexarelin is a synthetic hexapeptide agonist of the GHS-R1a, which is chemically more stable and functionally more potent when compared with ghrelin [35], which makes hexarelin a promising substitute for ghrelin [36]. Ghrelin has been demonstrated to protect several cell types such as adipocytes [37], osteoblast [38], cardiomyocytes and endothelial cells [39] by inhibiting apoptotic stimuli. As well, ghrelin has been shown to have protective effects *in vivo*, in rats exposes to status epilepticus [40] or in rat model of cisplatin-induced cachexia [4].

This study demonstrates anti-apoptotic effects of hexarelin via the inhibition of caspases-3 and -7 activation.

Neuro-2A cells treated for 24 h with increasing concentrations of H_2_O_2_ showed a significant activation of both caspases. Treatment of cells with hexarelin at 10^−6^ M attenuated the activation of caspase-3 and -7. Our hypothesis was that the modulation of caspase mRNA levels operated by hexarelin was dependent of the intracellular pro-apoptotic signalling molecules belonging to the BCL-2 family.

The BCL-2 family consist of two groups of mediators: the anti-apoptotic group, mainly represented by Bcl-2, and the pro-apoptotic group, principally Bax. Both groups play important roles in mitochondrial related apoptosis pathways [41].

Therefore, we quantified the effects of hexarelin on the modulation of the expression in Bcl-2 and Bax mRNA levels by RT-PCR.

As expected, H_2_O_2_ treatment of Neuro-2A cells induced the activation of pro-apoptotic Bax and the inhibition of anti-apoptotic Bcl-2, in a concentration dependent manner. Hexarelin treatment did not affect mRNA levels of apoptotic signalling molecules compared to the control group, demonstrating that hexarelin does not stimulate the apoptosis pathway. At the same time, the decrement of Bax mRNA levels and the increment in Bcl-2 mRNA expression, quantified by RT-PCR, in the group with 24 h of co-treatment confirmed the anti-apoptotic effect of hexarelin.

To investigate the molecular pathways involved in hexarelin neuroprotection, we investigated the expression of MAPKs (ERK and p38) and PI3K/Akt.

MAPKs activation contribute to neuronal dysfunction and are involved in NDDs [42,43]. Furthermore, ERK has been shown to participate in the regulation of cell growth and differentiation, response to cellular stress and cytokines [44].

In this study, treatment of Neuro-2A cells with H_2_O_2_ led to cell death by up-regulating p-ERK and p-p38 protein expression. The up-regulation of MAPKs induced by H_2_O_2_ stimulation were suppressed by hexarelin treatment. In addition, hexarelin alone didn’t modulate the MAPKs proteins level compare to the control.

PI3K/Akt is a key anti-apoptotic effector in the growth factor signalling pathway [45]. In particular, the phosphorylation of Thr-308 and Ser-473 of Akt serves a key role in mediating the anti-apoptotic actions of growth factors on cells and play an important role in neuronal protection [46,47].

In this study, H_2_O_2_-induced oxidative stress significantly increased the dephosphorylation of Akt which precipitated activation of the apoptotic pathway. Hexarelin treatment did not alter the p-Akt/t-Akt ratio compared to controls but in cells treated for 24 h with both hexarelin and H_2_O_2_, it significantly increased Akt protein expression compared with cells treated with H_2_O_2_ alone.

In conclusion, our findings demonstrate that H_2_O_2_ causes early and late apoptotic cell death in Neuro-2A cells. Treatment of cells with hexarelin suppresses cellular cytotoxicity, inhibits apoptosis and potentiates MAPKs and PI3K/Akt survival pathways.

This study demonstrates that hexarelin is able to protect Neuro-2A cells from H_2_O_2_-caused cytotoxicity effects but further experiments are needed to clarify the molecular mechanism of action, and if its effects are mediated by ghrelin receptor (GHS-R1a).

## Acknowledgments

RM, EB and AT conceived and designed the in vitro experiments. RM, LR, LM and EB performed the research, analysed the data and prepared figures. RM prepared the manuscript. AT, RJO, SC and VL supervised the project and revised the final manuscript. All of the authors have read and approved the manuscript.

